# Campycins are novel broad spectrum antibacterials killing *Campylobacter jejuni* by targeting the essential major outer membrane protein (MOMP)

**DOI:** 10.1101/2023.12.21.572796

**Authors:** Athina Zampara, Yilmaz Emre Gencay, Lone Brøndsted, Martine Camilla Holst Sørensen

**Author notes:** Correspondence: Martine Camilla Holst Sørensen. SNIPR Biome, Lersø Parkallé 44, København, 2100, Denmark (Yilmaz Emre Gencay).

## Abstract

Pyocins are high molecular weight bacteriocins produced by *Pseudomonas aeruginosa* that can be retargeted to new bacterial species by exchanging the pyocin tail fibers with bacteriophage receptor binding proteins (RBPs). Here, we develop retargeted pyocins called campycins as new antibacterials to specifically and effectively kill the major foodborne pathogen *Campylobacter jejuni.* We used two diverse RBPs (H-fibers) encoded by CJIE1 prophages found in the genomes of *C. jejuni* strains CAMSA2147 and RM1221 to construct Campycin 1 and Campycin 2, respectively. Together Campycin 1 and 2 could target all *C. jejuni* strains tested due to complementary antibacterial spectrums. In addition, both campycins led to more than 3 log reductions in *C. jejuni* counts under microaerobic conditions at 42°C, whereas the killing efficiency was less efficient under anaerobic conditions at 5°C. We furthermore discovered that both H-fibers used to construct the campycins bind to the essential major outer membrane protein (MOMP) present in all *C. jejuni,* in a strain specific manner. Protein sequence alignment and structural modelling suggest that the highly variable extracellular loops of MOMP form the binding sites of the diverse H-fibers. Further *in silico* analyses of 5000 MOMP sequences suggest that the protein fall into three major clades predicted to be targeted by either Campycin 1 or Campycin 2. Thus, campycins are promising antibacterials against *C. jejuni* expected to broadly target numerous strains of this human pathogen found in nature and agriculture.

**IMPORTANCE:** *Campylobacter jejuni* is the leading cause of bacterial foodborne gastroenteritis and responsible for more than 800 million cases globally each year posing a continuous risk to human health and a huge economic and societal burden. Here, we developed re-targeted R2 pyocins (campycins) as novel antibacterials against *C. jejuni* by using the receptor binding proteins of CJIE1 prophages observed in many *C. jejuni* genomes. Notably, campycins broadly target the highly variable yet essential major outer membrane protein (MOMP), and result in more than 3-log reductions in *C. jejuni* counts under conditions promoting bacterial growth. We therefore propose that campycins have the potential to lower *C. jejuni* colonization levels in the chicken gut, the main reservoir and cause of human disease, representing a novel efficient antibacterial solution specifically developed to target this widespread foodborne pathogen.

## INTRODUCTION

Pyocins are high molecular weight bacteriocins produced by *Pseudomonas aeruginosa* that resemble bacteriophage (phage) tail structures (1). Specifically, R-type pyocins look like non-flexible contractile phage tails related to the simple myovirus Enterobacteria phage P2 (2). R-type pyocins function as nano-needles that contract after binding to a bacterial surface receptor, thereby forming pores in the cell membrane. This results in the dissipation of the bacterial membrane potential leading to cell death. Several studies show that the bactericidal effect of R2-type pyocins can be redirected to new bacterial species other than *P. aeruginosa* by exchanging the original pyocin tail fibers with phage receptor binding protein (RBPs). Phage RBPs are highly specific in recognising and binding to surface components of their bacterial hosts and have therefore been exploited for designing several novel antibacterials (3,4). Examples are the receptor binding domains of RBPs from phage phiV10 infecting *Escherichia coli* O157:H7 and phage L-413c infecting *Yersinia pestis* that were fused to the N-terminal of the native R2 pyocin tail fibres resulting in chimeric R-type pyocins killing these new bacterial species (1,5). Thus, engineered R-type pyocins are promising as novel antibacterials targeting pathogenic and antibiotic resistant bacteria.

*Campylobacter jejuni* is one of the most common causes of human gastroenteritis worldwide. *C. jejuni* colonizes the poultry gut as a commensal and the main risk of human infection is consumption of raw or undercooked chicken meat (6,7). There are more than 800 million human cases of campylobacteriosis worldwide annually and the economic burden has been estimated to reach $1.56 billion in healthcare costs in the USA, $80 million in Canada, and €2.4 billion in the European Union annually (8). Thus, there is an urgent need for developing novel antibacterials to lower the prevalence of this pathogen either in the chicken gut or on the chicken meat. We recently succeeded in retargeting a R-type pyocin to *C. jejuni* using the tail fiber protein H (H-fiber) of a CJIE1 prophage (9,10). The retargeted pyocin, called Campycin 1, was able to bind the *C. jejuni* strain from which it was derived, however the killing efficiency and potential to target other *C. jejuni* strains were not determined. Also, the receptor recognized by the H-fiber remains to be identified, although Campycin 1 could bind *C. jejuni* independently of capsular polysaccharides and flagella recognized by the majority of virulent *C. jejuni* phages (10,11). The CJIE1 prophage was originally identified as an insertion element in *C. jejuni* RM1221 and has since been discovered in many *C. jejuni* genomes mostly as a cryptic prophage (9,12). While most of the CJIE1 genes across different *C. jejuni* strains are conserved, the H-fiber gene show significant sequence divergence, suggesting that different H-fibers may bind to different receptors on the *C. jejuni* surface (9). We recently found that virulent *Campylobacter* phage F341 encodes a novel RBP with a receptor binding domain demonstrating similarity to the H-fiber of a CJIE1 prophage in *C. jejuni* strain H5-4A-4 (13). While phage F431 is dependent on lipooligosaccharides for infection, the F341_RBP show very low sequence similarity to the H-fiber used to construct Campycin 1, suggesting that Campycin 1 may interact with a yet unknown receptor.

Here we developed novel antibacterials, campycins, against *Campylobacter* by re-targeting R2 pyocins using diverse RBPs (H-fibers) encoded by CJIE1 prophages. We further aimed to determine the receptors recognized by these H-fibers to predict strain sensitivity to campycins and to investigate the application potential in relation to broilers and chicken meat. We found that Campycin 1 and 2 demonstrated broad and complementary spectrums on diverse *C. jejuni* strains. While both campycins display bactericidal activities under conditions resembling the chicken gut, the natural niche and main reservoir of *C. jejuni*, their activities were significantly reduced under food storage conditions. We furthermore discovered that the highly variable, yet essential major outer membrane protein (MOMP) is the receptor for both H-fibers used for the campycins constructs and that their different antibacterial spectrums correlate with MOMP diversity. Finally, bioinformatic analysis of MOMP variants predicts that campycins could target a wide spectra of *C. jejuni* in nature thus showing great potential as effective antibacterials against this pathogen.

## RESULTS

### Exploiting the H-fiber diversity to construct campycins

To construct campycins with diverse killing spectrum, we aimed to use distinct H-fibers of CJIE1 prophages for re-targeting R2-pyocins killing a broad range of *C. jejuni* strains. Thus, we identified *C. jejuni* strains in our collection encoding CJIE1 prophages by analyzing their genome sequences. Our analysis showed that the CJIE1 prophage was present in five out of 15 sequenced *C. jejuni* CAMSA strains of Danish origin as well as the highly characterized lab strain *C. jejuni* RM1221. While all H-fibers originating from the CAMSA strains were highly conserved (Fig. S1), RM1221 encoded an H-fiber displaying significant sequence variability at the C-terminus (235-359 amino acids) (Fig. 1A). Since we previously constructed Campycin 1 using the H-fiber derived from CJIE1 of *C. jejuni* CAMSA2147 (10), the H-fiber from RM1221 was used to construct Campycin 2, expected to demonstrate a different antibacterial range than Campycin 1. To construct Campycin 2, we exchanged the receptor binding domain of the native tail fiber of the R2-pyocin in *P. aeruginosa* PAO1 with the C-terminus of the H-fiber gene of the CJIE1 prophage in RM1221 (last 209 amino acids) as done previously with Campycin 1 (Fig. 1B). Like CAMSA2147, the gene downstream of the RM1221 H-fiber gene encodes a protein that harbors a DUF4376 domain also found in the RBP chaperone of the phage Mu (14,15). This putative chaperone is highly conserved between the CAMSA2147 and RM1221 derived CJIE1 prophages (Fig. S2). We previously showed that Campycin 1 was only able to bind and cause lysis on a CAMSA2147 lawn when the putative chaperone was co-expressed (10). We therefore also included the RM1221-derived CJIE1 putative chaperone for expressing Campycin 2 to investigate whether it is required for activity.

**FIG 1.**
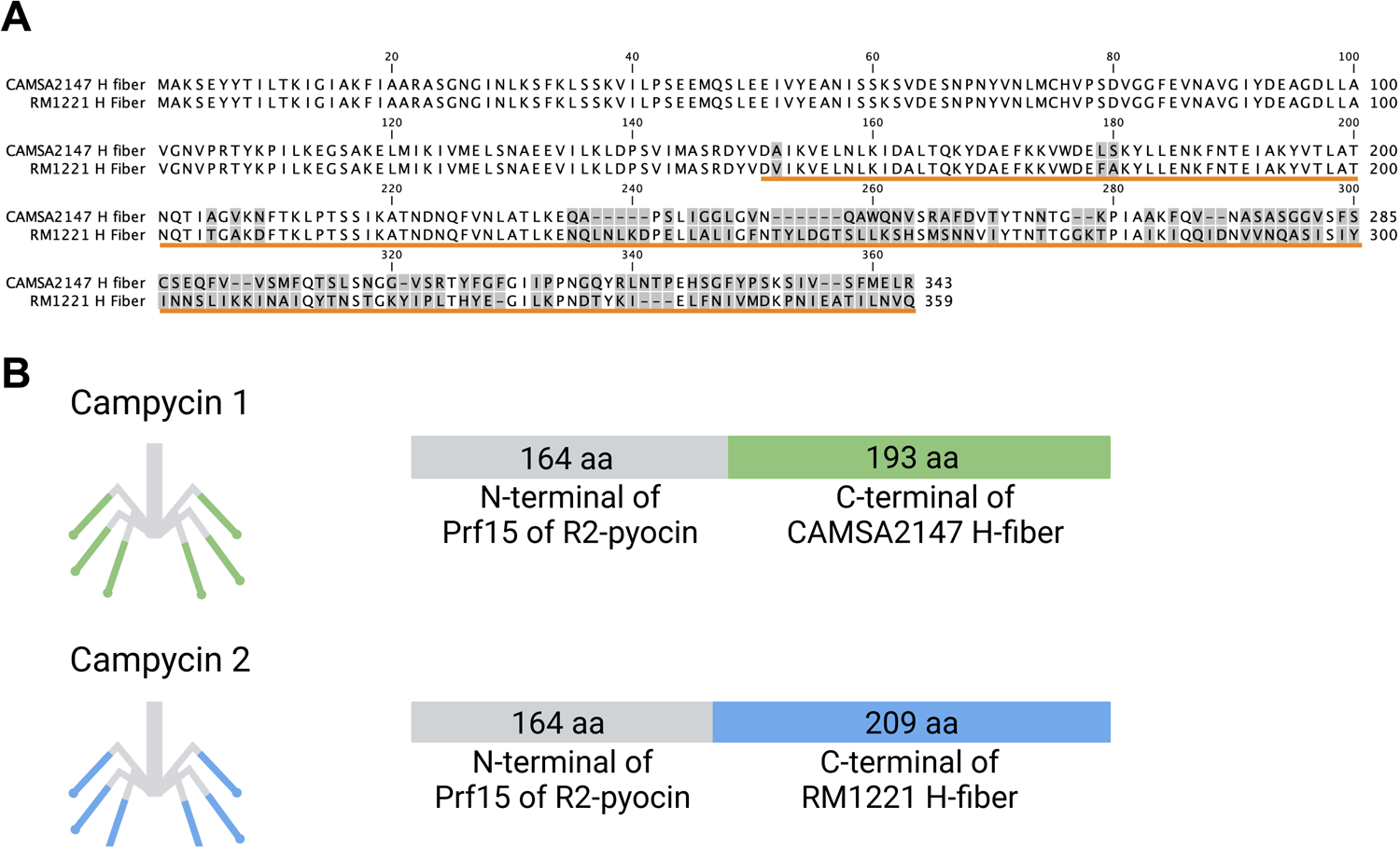
H-fibers from CJIE1 prophages in *C. jejuni* strains CAMSA2147 and RM1221 and the construction of campycins. (**A**) Alignment of H-fibers from CJIE1 prophages present in CAMSA2147 (SAMN08987259) and RM1221 (SAMN14933898). Protein sequences were aligned using CLC Main Workbench 22 (QIAGEN) and divergent amino acids are indicated by grey boxes. The orange line indicates the amino acids present in Campycin 1 and 2. (**B**) Schematic representation of Campycin 1 and 2. Campycin 1 contains 164 amino acids of the N-terminus of Prf15 tail fiber originating from *P*. aeruginosa PAO1 R2-pyocin (grey) fused to 193 amino acids of the C-terminus of the H-fiber derived from the CAMSA2147 CJIE1 prophage (green) (10). Campycin 2 contains the N-terminus Prf15 encoded *P. aeruginosa* PAO1 R2-pyocin (grey) fused to 209 amino acids of the C-terminus of the H-fiber derived from the RM1221 CJIE1 prophage (blue).

### Campycins show broad antibacterial spectrum targeting diverse *C. jejuni* strains

To identify the activity spectrum of Campycin 1 and 2, purified campycins co-expressed with or without the downstream H-fiber chaperones were spotted on bacterial lawns of a collection of *C. jejuni* strains. A clear zone on the bacterial lawn indicates that the campycin binds and thus kills the cells. As a control, we spotted the native R2 pyocin on the *C. jejuni* lawns and as expected no clearing zones were observed. Campycin 1 killed 29 out of the 36 strains, revealing a broad spectrum, but only when co-expressed with its chaperone (Fig. 2A). Campycin 2 killed the remaining 7 strains in our collection that were not sensitive to Campycin 1, while it did not kill the 29 strains that were sensitive to Campycin 1 thereby showing a complementary coverage. Interestingly, in contrast to Campycin 1, the predicted chaperone was not required for Campycin 2 to exert activity. These results confirm that the H-fiber sequence diversity promoted distinct antibacterial spectrum of the two campycins. Previously, antimicrobial resistance of *Campylobacter* has been correlated with multilocus sequence types (MLST) (16,17). However, the sensitivity to either Campycin 1 or Campycin 2 showed no correlation with the assigned MLST of the *C. jejuni* strains tested (Fig. 2A). In conclusion, Campycin 1 and 2 displayed complementary antibacterial activities, thus proving that the natural diversity of *C. jejuni* CJIE1 H-fiber leads to recognition of diverse host receptors.

**FIG 2.**
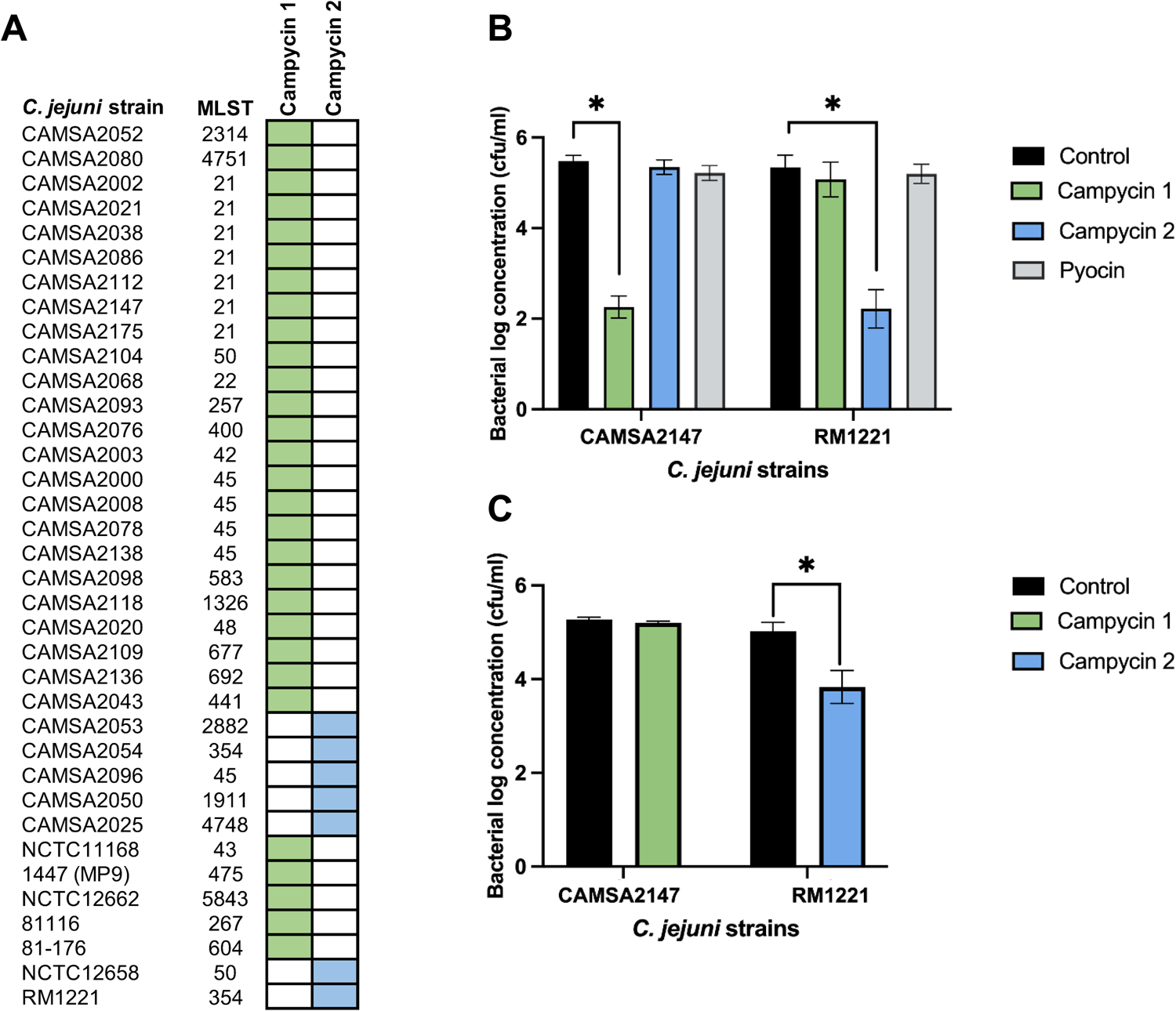
Antibacterial spectrum of Campycin 1 and 2. **(A**) Antibacterial coverage of campycins. Purified campycins were spotted on bacterial lawns of each of the *C. jejuni* strains listed. Campycin 1 was co-expressed with the downstream H-fiber chaperone, whereas Campycin 2 was expressed both with and without the H-fiber chaperone showing the same results. MLST: Multilocus sequence type. Green and blue: clearing zone on the bacterial lawn for Campycin 1 and Campycin 2, respectively. White: no clearing zone observed. (**B**) Antibacterial activity of campycins after incubation for 3 hours at microaerobic conditions at 42 °C. 100 μl of each campycin (10^9^ particles/ml) was mixed with 100 μl of each *C. jejuni* strain (10^6^ cfu/ml). As controls TN50 buffer or native R2 pyocins were used. Log reductions represent cfu/ml compared to the TN50 buffer control and error bars are the standard deviations of the mean from biological triplicates. *Significant reduction at *P* < 0.05. (**C**) Antibacterial activity of campycins after incubation for 24 hours at 5 °C under anaerobic conditions. 100 μl of each campycin (10^9^ particles/ml) was mixed with 100 μl of each *C. jejuni* strain (10^6^ cfu/ml). As controls TN50 buffer and native R2 pyocin were used. Log reductions represent cfu/ml compared to the TN50 buffer control and error bars are the standard deviations of the mean from biological triplicates. Significant reduction at *P* < 0.05.

### Campycin 1 and 2 exert efficient antibacterial activity

To assess the killing efficiency of Campycin 1 and 2, *C. jejuni* CAMSA2147 and RM1221 cells were treated with either of the campycins or the native R2 pyocin under different conditions related to the chicken gut and chicken meat. Campycin 1 led to 3,2± 0,2 log reduction of CAMSA2147 after 3 hours of co-incubation at 42 °C under microaerobic conditions resembling conditions in the chicken gut (Fig. 2B). Similarly, Campycin 2 killed RM1221 leading to 3,1 ± 0,5 log reduction of cells under these conditions. No significant reduction was noticed in any of the strains after R2 pyocin application under the same conditions (Fig. 2B). Interestingly, no colonies were recovered after 24 hours at 42 °C under microaerobic conditions when CAMSA2147 or RM1221 were treated with Campycin 1 or Campycin 2, respectively, suggesting that killing is ongoing, and a higher log reduction is achieved over time. To evaluate the use of campycins as antibacterials in refrigerated food, we tested their killing efficiencies at 5 °C under anaerobic conditions after 24 hours of incubation. Campycin 1 led to only 0,1 ± 0,1 log reduction of CAMSA2147 and Campycin 2 to 1,2 ± 0,6 log reduction of RM1221 (Fig. 2C). Thus, antibacterial activities of campycins are more efficient under microaerobic conditions promoting bacterial growth related to the avian gut compared to food packaging conditions.

### The major outer membrane protein (MOMP) is the receptor of the H-fibers used to construct Campycin 1 and 2

To identify the surface receptors targeted by Campycin 1 and 2, we conducted a pull-down assay using purified C-terminal domains of the CAMSA2147 and RM1221 CJIE1 H-fibers (H-fiber_C-term_) co-expressed with their downstream chaperone as bait and cell lysates of CAMSA2147 and RM1221 as prey (Fig. 3). When purified CAMSA2147 H-fiber_C-term_ was ran on the gel, a band at approximately 20 kDa could be observed (Fig. 3A). Further analysis using mass spectrometry revealed that the band was a mixture of the CAMSA2147 H-fiber_C-term_ (22 kDa) and the downstream chaperone (20 kDa) demonstrating that chaperone is co-eluted with the H-fiber, thus indicating that the chaperone binds to the H-fiber (Table S5). When the CAMSA2147 H-fiber_C-term_ was pre-incubated with a CAMSA2147 cell lysate, an additional band was detected at approximately 50 kDa, demonstrating that CAMSA2147 H-fiber binds to a protein of this size (Fig. 3A). The interating protein was identified as the essential major outer membrane protein (MOMP) which is porin of 46 kDa encoded by the *porA* gene (18) (Table S5). Thus, MOMP encoded by CAMSA2147 is the receptor of the CAMSA2147 H-fiber. As expected, no additional band was observed when the CAMSA2147 H-fiber_C-term_ was incubated with a RM1221 cell lysate (Fig. 3A), supporting that RM1221 does not encode the receptor recognized by the CAMSA2147 H-fiber, thus explaning why Campycin 1 is not able to kill *C. jejuni* RM1221.

**FIG 3.**
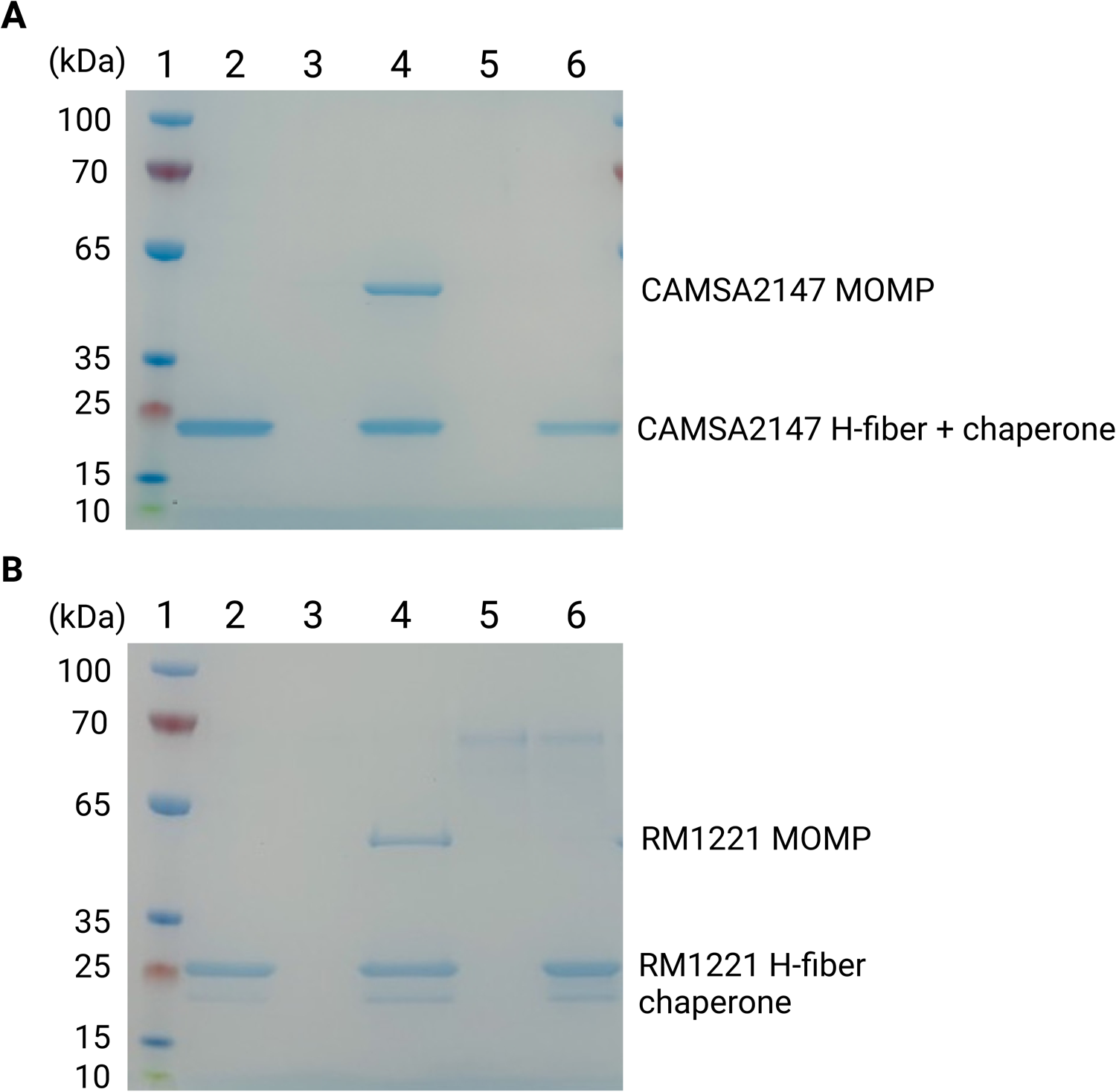
SDS-PAGE of pull-down assays with purified H-fibers and cell lysates from *C. jejuni* CAMSA2147 and RM1221. The C-terminus of the CAMSA2147 or RM1221 CJIE1 H-fibers were fused to a His-tag and co-expressed with the downstream chaperones. Purified H-fibers were used as bait in pull-down assays and CAMSA2147 or RM1221 cell lysates as prey. The eluted fractions were analysed by SDS-page. (**A**) Pull-down assay with purified CAMSA2147 H-fiber. Lane 1: marker, lane 2: CAMSA2147 H-fiber co-expressed with the chaperone, lane 3: CAMSA2147 cell lysate, lane 4: CAMSA2147 cell lysate incubated with CAMSA2147 H-fiber co-expressed with the chaperone, lane 5: RM1221 cell lysate, lane 6: RM1221 cell lysate incubated with CAMSA2147 H-fiber co-expressed with the chaperone. (**B**) Pull-down assay with purified RM1221 H-fiber. Lane 1: marker, lane 2: RM1221 H-fiber co-expressed with the chaperone, lane 3: RM1221 cell lysate, lane 4: RM1221 cell lysate incubated with RM1221 H-fiber co-expressed with the chaperone, lane 5: CAMSA2147 cell lysates, lane 6: CAMSA2147 cell lysates incubated with RM1221 H-fiber co-expressed with the chaperone.

Similarly, to elucidate the receptor of the RM1221 CJIE1 H-fiber, we analysed the purified RM1221 H-fiber_C-term_ co-expressed with the downstream chaperone with or without pre-incubation using a RM1221 cell lysate (Fig. 3B). Our results showed two bands at approximately 20 and 25 kDa when the purified RM1221 H-fiber_C-term_ was ran on the gel (Fig. 3B). These bands were confirmed to be the RM1221 H-fiber_C-term_ (24 kDa) and the downstream chaperone (20 kDa), demonstrating that also in this case the chaperone interacts and co-elutes with the H-fiber (Table S5). When the purified RM1221 H-fiber_C-term_ was incubated with a RM1221 cell lysate, an additional band at approximately 50 kDa was visible, demonstrating that the RM1221 H-fiber binds to a protein of this size. Interestingly, this band was again identified as MOMP (47 kDa) demonstrating that the RM1221 H-fiber also use MOMP in RM1221 as a surface receptor (Table S5). Yet, an interacting protein was not observed on the gel when the purified RM1221 H-fiber_C-term_ was incubated with a CAMSA2147 cell lysate (Fig. 3B), demonstrating that the RM1221 H-fiber does not bind to MOMP encoded by CAMSA2147.

Overall, our results revealed that the H-fibers of the CJIE1 prophages in CAMSA2147 and RM1221 both use MOMP as a receptor yet in a strain-specific manner, explaining the different spectrums of the two campycins.

### Diversity of MOMP in *C. jejuni* corelates with the antibacterial spectra of campycins

To investigate the strain specificity of the H-fibers derived from CAMSA2147 and RM1221, we compared the MOMP protein sequences of the two strains. MOMP is a trimer of 18-stranded antiparallel β-barrel monomers with eight extracellular surface-exposed loops out of which the four (L1, L3, L4, and L6) fold inside of the barrel (Fig. 4A) (18). Particularly the loop regions demonstrate large sequence diversity among *C. jejuni* strains (19). Our results showed that the CAMSA2147 and RM1221 MOMPs displayed significant variability in multiple positions, but mostly in the loop regions (Fig. 4B). To further investigate if the sequence variability observed in the MOMP loop regions correlate with the sensitivity of the *C. jejuni* strains towards the two different campycins, we conducted an *in-silico* comparison of MOMP homologs of the strains tested for campycin sensitivity. As expected, the MOMP protein sequences of the strains sensitive to Campycin 1 were highly similar across the entire sequence, and likewise for the strains sensitive to Campycin 2 (Fig. 4C). Thus, the specificity of the CAMSA2147 and RM1221 H-fibers correlates with MOMP diversity particularly in the extracellular surface-exposed loops, explaining the different target spectrums of the two campycins.

**FIG 4.**
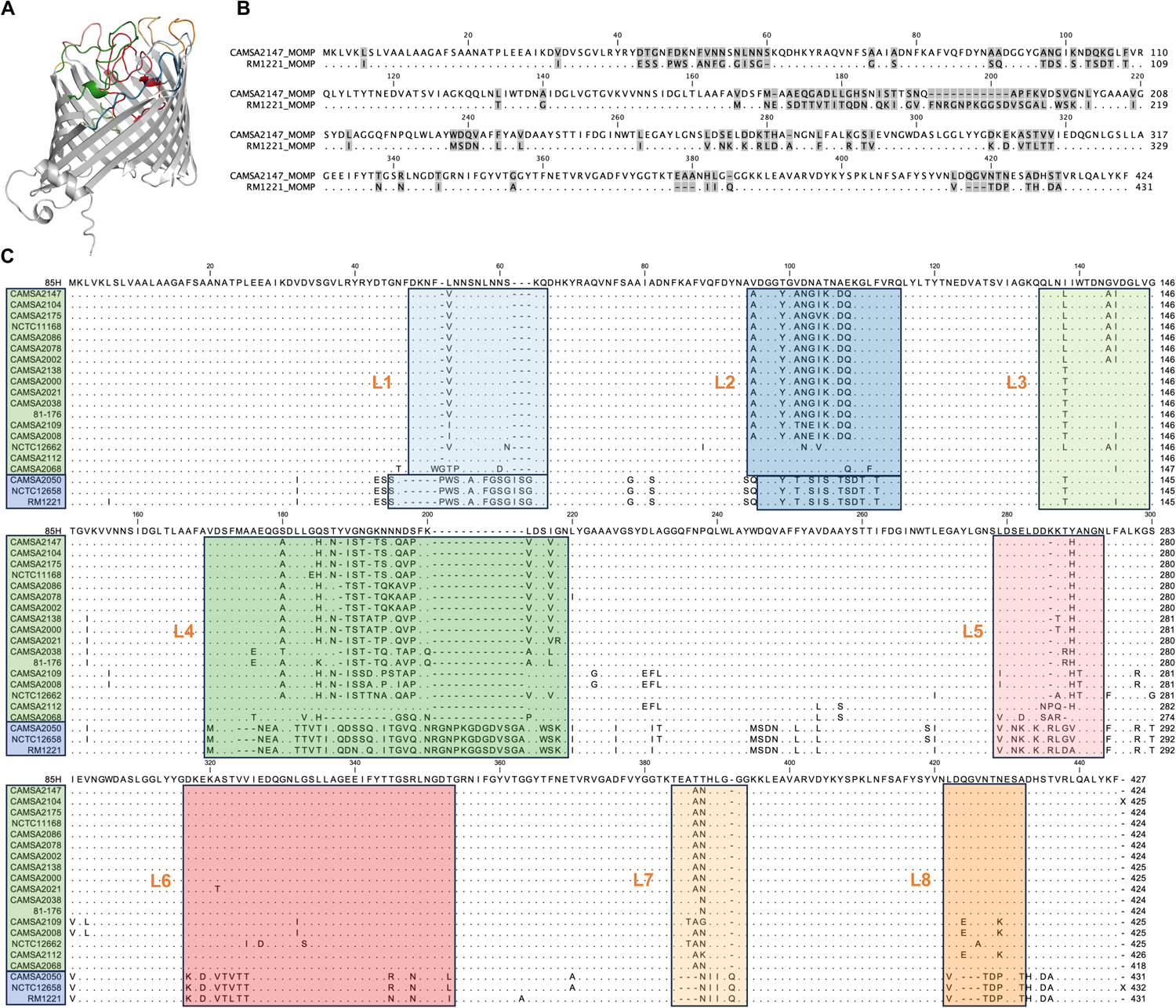
Alignment of MOMP homologs from *C. jejuni* strains sensitive to Campycin 1 and 2. **(A**) MOMP monomer from *C. jejuni* CAMSA2147 as predicted by AlphaFold2. The extracellular surface-exposed loops are indicated by color corresponding to colors used in the MOMP protein sequence alignment in (C). Model confidence score is listed in Table S4. (**B**) Protein sequence alignment of MOMP from *C. jejuni* CAMSA2147 and RM1221. Grey boxes indicated divergent amino acids. (**C**) MOMP protein sequence alignment of *C. jejuni* strains tested for sensitivity towards Campycin 1 and 2, respectively. MOMP from *C. jejuni* 85H (5LDV) is added as a reference and conserved amino acids are indicated as dots. Strains sensitive to Campycin 1 are grouped in a green box and strains sensitive to Campycin 2 are grouped in a blue box on the left-hand side. Extracellular surface-exposed loops are indicated by boxes of different colors and listed as L1-L8. Protein sequences were aligned using CLC Main Workbench 22 (QIAGEN).

### Structural prediction of CJIE1 H-fibers and H-fiber-MOMP complex suggests interaction at multiple extracellular surface-exposed loops of MOMP

To further explore the interaction between the CJIE1 H-fibers and MOMP we used Alphafold and Alphafold multimer for predicting the H-fiber structures and the H-fiber-MOMP complexes (Fig. 5). Structural prediction of the CAMSA2147 and RM1221 CJIE1 H-fibers as homotrimers showed that both displays typical tail fiber morphology i.e., long and slim with N-terminal globular shoulder domains, shafts/stems formed primarily by α-helices and a distal C-terminal knob-like structure consisting of antiparallel ß-sheets forming distal loops likely interacting with the receptor. The H-fiber C-terminus used in the campycin constructs (151-343 aa for CAMSA2147 H-fiber and 151-359 aa for RM1221 H-fiber) comprise the entire stem as well as the distal knob (Fig. 5A). Residues 235-343/359 demonstrating significant dissimilarity between the two H-fibers form the distal knob thus the receptor binding region. Interestingly, the knob of the RM1221 H-fiber is slightly wider compared to the H-fiber of CAMSA2147, suggesting also minor structural difference between the two H-fibers (Fig. 5A). The model of the CAMSA2147 H-fiber-MOMP complex predicts that most of the MOMP extracellular surface-exposed loops (except L3 and L8) interact with the CAMSA2147 H-fiber (Fig. 5B). The same extracellular loops were predicted to interact with the H-fiber when modelling the H-fiber-MOMP complex from RM1221 as well (data not shown). These loops are also located in the region showing most sequence divergence when comparing MOMPs of CAMSA2147 and RM1221 (Fig. 5D). Furthermore, the model of the CAMSA2147 H-fiber-MOMP complex predicts that it is the three distal loops of the H-fiber knob domain that are responsible for interacting with the MOMP receptor (Fig. 5B and C). These three loops are formed by amino acids 278-285, 298-304 and 323-333 in CAMSA2147, whereas the distal loops of the RM1221 H-fiber are formed by amino acids 285-296, 310-325 and 341-350 (Fig. 5D). Thus, structural modelling confirms that the C-terminal region demonstrating the largest sequence diversity of the CAMSA2147 and RM1221 CJIE1 H-fibers compose the receptor binding domain. Furthermore, several highly variable extracellular loops of MOMP are predicted to form the receptor binding sites of these H-fibers.

**FIG 5.**
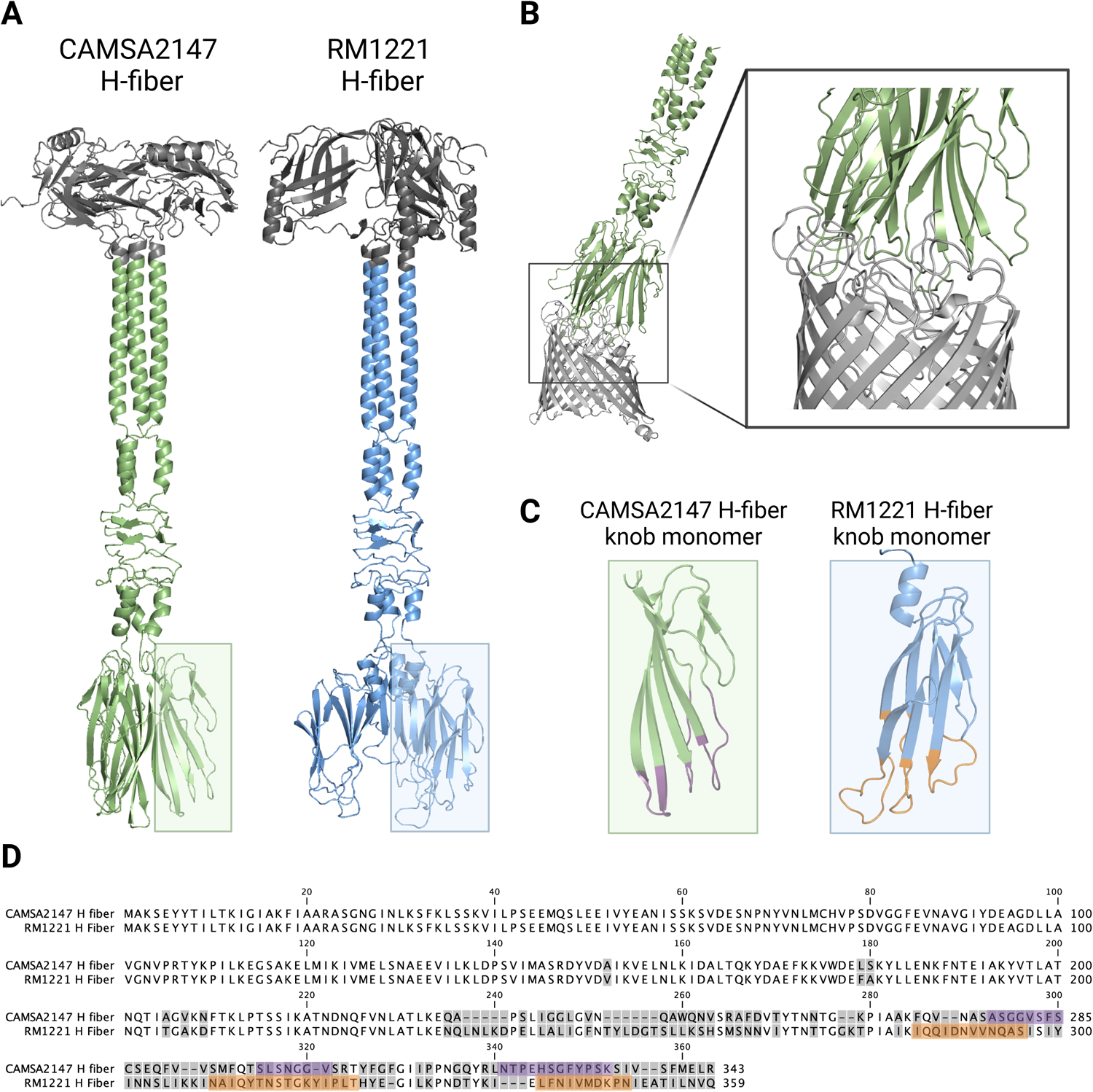
Structural prediction of the CAMSA2147 and RM1221 H-fibers and the CAMSA2147 H-fiber-MOMP complex. (**A**) H-fibers predicted as homotrimers. The green and blue colors, respectively, indicate the part of the H-fibers used in the campycin constructs. (**B**) AlphaFold2 model of the CAMSA2147 H-fiber-MOMP complex showing interaction between the three distal loops in the knob region of the H-fiber and the extracellular surface-exposed loops of MOMP. (**C**) and (**D**) The three distal loops (highlighted in purple and orange, respectively) in the knob of the CAMSA2147 and RM1221 H-fibers demonstrate significant sequence divergence. Protein sequences of the CAMSA2147 and RM1221 H-fibers were aligned using CLC Main Workbench 22 (QIAGEN). Divergent amino acids are highlighted in grey and found in the C-terminal region. All structural models were created using Colab AlphaFold2 and visualized using PyMOL. Model confidence scores are listed in Table S4.

### *C. jejuni* MOMPs form three major phylogenetic clades predicted to be targeted by either Campycin 1 or Campycin 2

The large sequence diversity of MOMP in different *C. jejuni* strains may be a potential drawback for the application of campycins in agriculture. Thus, to predict whether the campycins could target a wide range of diverse *C. jejuni*, we analyzed the diversity of *C. jejuni* MOMPs using sequences available in the public database NCBI. CAMSA2147 MOMP (WP_002875862.1) was used as a query to retrieve 5000 non-redundant MOMP homologs that were aligned and used for construction of an unrooted phylogenetic tree. The retrieved MOMP homologs grouped into three major clades called I, II and III (Fig. 6). Interestingly, MOMPs derived from *C. jejuni* strains in our collection are present within all distinct clades i.e., MOMP variants from strains targeted by Campycin 1 are present in clades II and III, while MOMP variants from strains targeted by Campycin 2 are present in clade I (Fig. 6). These results suggest that MOMP variants likely targeted by either Campycin 1 or Campycin 2 are widespread in *C. jejuni* promoting campycins as broad spectrum antibacterials against this pathogen.

**FIG 6.**
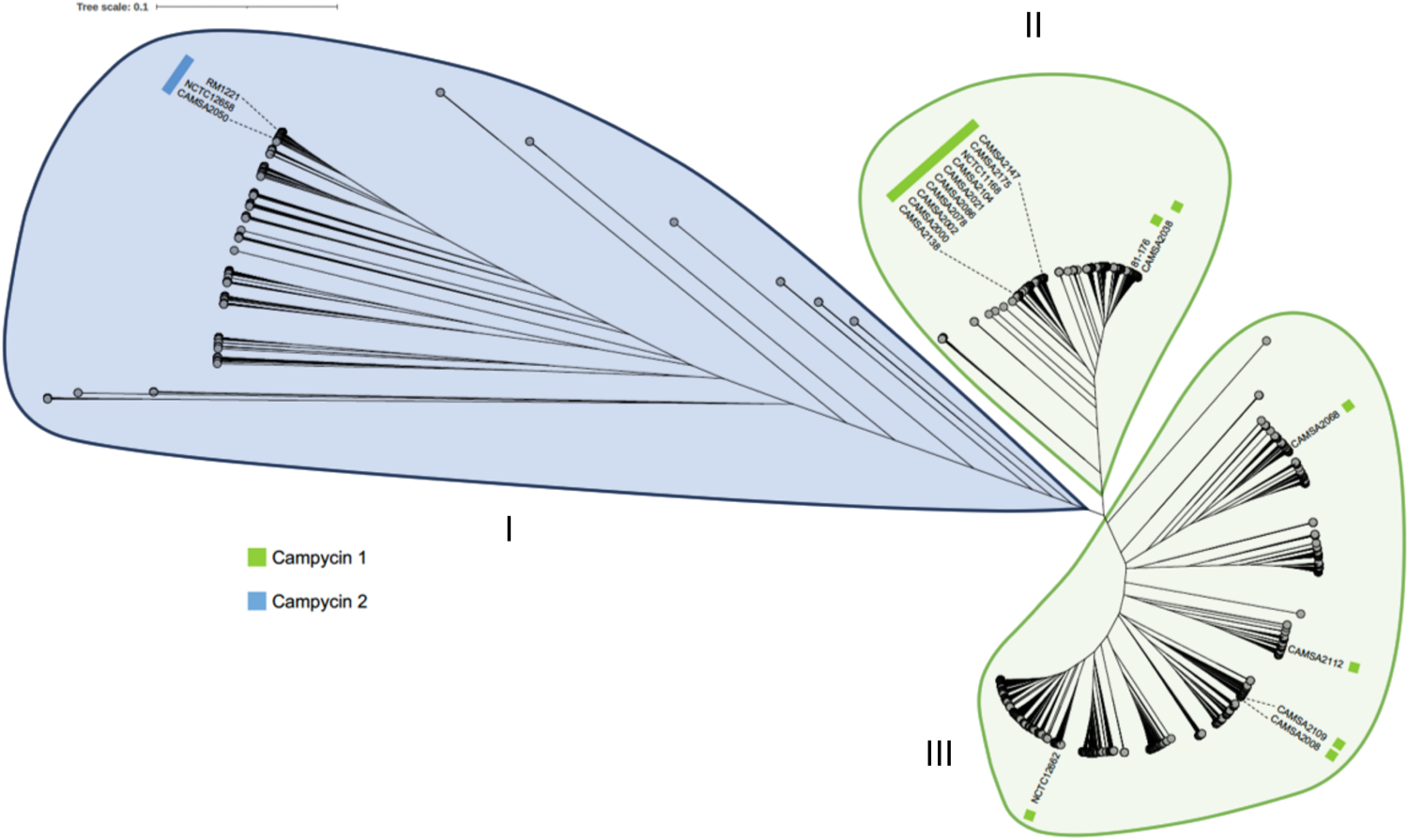
Phylogenetic tree of 5000 non-redundant MOMP variants showing three distinct clades (I, II and III). An unrooted phylogenetic tree was created by using 5000 *C. jejuni* MOMP sequences retrieved from the National Center for Biotechnology Information (NCBI) genome database using BLASTP (34) and MOMP from CAMSA2147 (WP_002875862.1). Sequences were aligned in mafft (35) and a neighbor-joining phylogenetic tree was created using 100 bootstraps. Display, manipulation and annotation of the phylogenetic tree was further conducted using Interactive Tree OfLife (iTOLv6) (36). Further labeling highlight MOMPs from the *C. jejuni* strains in our collection and their sensitivity to Campycin 1 (green box) or Campycin 2 (blue box).

## DISCUSSION

*Campylobacter jejuni* poses a serious threat to public health and currently there are no available antibacterial solutions to eliminate this pathogen through the food production chain (20). Here, we exploit the target specificity of phages and the killing efficiency of pyocins to develop novel antibacterials against *C. jejuni.* We used diverse tail fibers (H-fibers) of *C. jejuni* CJEI1 prophages to re-target R2 pyocins and developed Campycin 1 and 2 with broad and complementary killing spectrum. We furthermore showed that the H-fibers used for construction of the campycins bind to the major outer membrane protein (MOMP), which is essential for *Campylobacter* growth, making campycins promising antibacterials against this pathogen.

Phage receptor binding proteins (RBPs) are key determinants for the host range of a phage and demonstrate large diversity to match the variety of surface structures expressed on the bacterial cell (21). We used two distinct RBPs (H-fibers) encoded by the CJIE1 prophages in *C. jejuni* strains CAMSA2147 and RM1221 for development of Campycin 1 and Campycin 2, respectively. While Campycin 1 showed binding and hence killing activity only when co-expressed with its downstream chaperone, Campycin 2 functioned independently of chaperone co-expression. Interestingly, a pull-down assay demonstrated that both H-fibers used in the campycin constructs interact physically with their downstream chaperone as they co-eluted during recombinant H-fiber protein expression and purification. Similar results have been observed for the tail fibers of *Alteromonas* phages V22 and A5, where the V22 tail fiber chaperone is essential for production of active tail fibers, whereas phage A5 is able to produce functional fibers without chaperone co-expression. Yet, in both cases the respective chaperone is co-eluted after recombinant protein purification, demonstrating tail fiber-chaperone interactions (22). Based on predictive modelling the authors propose that the tail fiber chaperones have alternative functions as transient caps for temporal shielding of the tail fiber tips to prevent interaction with cell debris during phage progeny release. Indeed, using Alphafold a similar complex was predicted for both the CAMSA2147 and RM1221 H-fibers and their chaperones, suggesting a similar alternative function also for chaperones of the *C. jejuni* CJIE1 prophages (Fig. S3).

Using a pull-down assay, we identified the major outer membrane protein (MOMP) encoded by *porA* as the receptor for the H-fibers used for campycin construction. MOMP is a porin, composed of 18-stranded antiparallel β-barrel monomers with eight external surface-exposed loops, and is mainly responsible for nutrient transport and the transport of other small molecules (18). In contrast to other gram-negative bacteria like *E. coli* that encode two major porins, OmpC and OmpF, *C. jejuni* only encodes a single MOMP which is essential for bacterial growth and present in all characterized *C. jejuni* (18). Although the structure of MOMP is highly conserved, the surface-exposed loops demonstrate high sequence variation when comparing different *C. jejuni* strains (19). The variation in the *porA* allele was therefore also proposed as a typing method for epidemiological studies of *C. jejuni* (23,24). Comparing MOMP in CAMSA2147 and RM1221 demonstrated sequence variation in most of the extracellular loops which also correlated with all *C. jejuni* strains tested for campycin sensitivity i.e., strains targeted by either Campycin 1 or Campycin 2 show similar sequences in the MOMP loop regions. Using AlphaFold to model the CAMSA2147 H-fiber-MOMP complex showed that the distal loops of the H-fiber knob region are interacting with most of the extracellular loops of MOMP. Thus, the structural prediction supports that differences in the extracellular MOMP loop sequences are associated with different binding capacities of the two H-fibers. Yet, although artificial intelligence tools for modelling protein structures have greatly improved, predicting the interaction between loop regions that are quite flexible remains challenging. Also, modelling protein interactions of different species i.e., bacteria and phage, is difficult often resulting in models with low confidence scores. Thus, experimental data is needed to verify the interacting residual components of the H-fibers and MOMP.

A major obstacle for new antimicrobials is the application range and potential of resistance development. Although MOMP is an essential porin found in all *C. jejuni,* the variation in the extracellular loops will indeed impact the campycin sensitivity. Yet, our analysis of 5000 MOMP sequences showed that these MOMP fall into three major clades and each clade contain MOMP from *C. jejuni* strains in our collection sensitive to either Campycin 1 or Campycin 2. Thus, we expect that the majority of MOMP present in *C. jejuni* belong to either of these clades thus anticipated to be sensitive to either Campycin 1 or Campycin 2. This suggest that a large majority of *C. jejuni* strains occurring in nature and agriculture will be targeted and killed by the campycins. Furthermore, as the campycins are predicted to interact with multiple extracellular loops of MOMP, we foresee that resistance development may require multiple mutational events. Also, we propose that correlating sequences of the CJIE1 H-fiber and MOMP variant within a given *C. jejuni* strain, would promote the identification of new H-fiber variants that can be used to develop additional campycins targeting more MOMP variants.

For application purposes related to food storage conditions (5°C and anaerobic atmosphere) campycins reduced *C. jejuni* cells at maximum 1,2 ± 0,6 logs. Lower temperature generally affects the fluidity of the outer membrane that thereby becomes more rigid (25). We speculate whether this might prevent the campycins from forming proper pores in the membrane which would reduce the effectivity. Further studies are needed to clarify if this indeed could be the case. In contrast, both Campycin 1 and 2 demonstrated high bactericidal activity exceeding 3-log reduction after 3 hours of incubation under microaerobic conditions, while no cells were recovered after 24 hours. These conditions relate to the main niche of *C. jejuni,* the avian gut, where particularly broiler colonization is a major cause of human disease (26). It is estimated that reducing the number of *C. jejuni* in the chicken gut by 3 log will lead to 58% less human cases of campylobacteriosis associated with consumption of chicken meat (27). *In vivo* studies using mice and rabbit models have shown that re-targeted pyocins are able to prevent gut colonization and symptoms of enteric disease development of *Clostridium difficile* and *E. coli* O157:H7 when delivered orally (28,29). Thus, addition of campycins to the drinking water for broiler chickens may be an applicable and easy approach to lower *C. jejuni* colonization levels in broilers thereby reducing the number of human cases. Consequently, *in vivo* studies are needed to evaluate the efficiency, safety, and application perspective of campycins in terms of use in the broiler production.

In conclusion, the campycins are promising novel antibacterials against *C. jejuni* predicted to target multiple strains present in agriculture.

## MATERIALS AND METHODS

### Bacterial strains

Bacterial strains used in this study are listed in Table S1. Tryptic Soy Broth (TSB) and Tryptic Soy Agar (TSA) were used for growing *P. aeruginosa*, while 15 µg/ml gentamicin was added for selection of *P. aeruginosa* transformants. *C. jejuni* strains were routinely grown on Blood Agar Base II (Oxoid) supplemented with 5% calf blood (BA) at 42 °C under microaerobic conditions (MA: 6% CO_2_, 6% O_2_, 84.5% N_2_ and 3.5% H_2_). Luria-Bertani broth (LB) and LB agar (LA) (Difco) were used for growing *E. coli*, while *E. coli* transformants were selected in the presence of 100 μg/ml kanamycin and 50 μg/ml chloramphenicol.

### Construction of campycins

Plasmids responsible for Campycin 2 expression were constructed similar to previous work (10) (Table S2). Plasmid pM50, previously engineered to carry the coding sequence of the N-terminal domain of R2-pyocin Prf15 (amino acids 1 to 164), was used as a backbone for inserting the RM1221 CJIE1 prophage H-fiber (CJE0231) along with or without the predicted chaperone (CJE0230). Specifically, pM50 was linearized via inverse PCR using R2N primers (Table S3) and the fragment comprising the coding sequence of the C-terminus of the H-fiber (amino acids 151 to 359) and the downstream predicted chaperone gene were amplified from RM1221 (CP000025) by specific primers (Table S3), adding overhangs identical to the distal ends of the linearized pM50. In-Fusion^®^ cloning – Takara Bio kit was used for the recombination, creating plasmids pM179 and pM182, respectively (Table S2). *P. aeruginosa* PAO1 Δ*prf15* was transformed with either of the plasmids and transformants were selected on TSA plates supplemented with 15 µg/ml gentamicin as described previously (30). As a result, a tail fiber deficient R2-pyocin derivative (Δ*prf15*) substituted *in trans* with the RM1221 C-terminal H-fiber along with or without the predicted chaperone.

### Campycin expression and purification

Expression and purification of R2-pyocin and campycins were conducted as previously described (1,10). Briefly, *P. aeruginosa* PAO1 expressing the native R2-pyocin or campycins were grown overnight in TSB at 37 °C and 100-fold diluted in G medium (31), that was supplemented with 15 µg/ml gentamicin in case of campycins. Cultures were incubated at 37 °C until reaching an OD_600_ of 0.25 and treated with mitomycin (3 µg/ml) and isopropyl-beta-D-thiogalactopyranoside (0.25 mM) for induction of expression. After 2.5 h of incubation at the same conditions, DNase I (Invitrogen) was added at a final concentration of 5 U/ml and the culture was further incubated for 30 min at the same conditions. Centrifugation of cultures (at 18,000 *g* at 4 °C for 1 h) was followed to remove cell debris and supernatants were treated with saturated ammonium sulfate solution (final concentration of 1.6 M), while stirring on ice. After overnight incubation at 4 °C, suspensions were centrifuged (18,000 *g* for 1 at 4 °C) and pellets were resuspended in 1/10^th^ of the start volume with ice cold TN50 buffer (50 mM NaCl and 10 mM Tris-HCl adjusted to pH 7.5). Ultracentrifuge was followed for 1 h at 60.000 g and pellets containing the precipitated campycins/pyocins were harvested and resuspended in 1/20^th^ of the start volume with ice cold TN50 buffer.

### Campycin quantification

Quantitative campycin assays were performed by counting surviving bacteria, using a method slightly modified from that described by Kageyama et al (32). Accordingly, CAMSA2147 or RM1221 were used for identifying the concentration of Campycin 1 (co-expressed with downstream chaperone) or Campycin 2, respectively. Cells were harvested by using 3 ml calcium Brain-Heart Infusion Broth (CBHI) and adjusted the optical density OD_600_ to 0.2 that equals approximately 10^9^ colony forming units per milliliter (cfu/ml). Campycins were fivefold diluted in TN50 buffer and 100 μl of each diluted campycin was mixed with 100 μl of adjusted *C. jejuni* in a deep well plate and incubated for 3 h at 42°C under MA conditions. TN50 buffer was used as a negative control. Proper dilutions were made in TN50, and samples were spotted on BA plates that were incubated under MA conditions at 42 °C. It was previously estimated for pyocins that a microtiter well that has an average of 1 pyocin particle per bacterium will yield at or near equilibrium 37% survivors, and a well with an average of 2.3 pyocin particles per bacterium will yield 10% survivors (5). Since campycins are engineered pyocins, we used this calculation to quantify campycins as well. A typical density of purified, concentrated campycins was 1 × 10^9^ per ml.

### Antibacterial spectrum of campycins

To determine the spectrum of campycins, a lysis spot assay was performed onto *Campylobacter* bacterial lawns as previously described (33). Briefly, cells were streaked on BA plates and harvested with calcium Brain-Heart Infusion Broth (CBHI). Suspensions were further adjusted to an OD_600_ of 0.35 and incubated for 4 h at 42 °C under MA conditions. Cell cultures (500 μl) were mixed with molten NZCYM overlay agar (5 ml of NZCYM broth [Sigma] with 0.6% agar [Sigma]) and poured on NZCYM basal agar (with 1.2% agar [Sigma]) plates, supplemented with 10 μg/ml vancomycin [Sigma]. After 20 min, plates were dried for 45 min in the flow hood and campycins (5 μl) were spotted on top, followed by incubation for 24 h at 42 °C under MA conditions. As negative controls, the native R2-pyocin or fiber mutant derivative (Δ*prf15*) were also spotted. Bacterial lawns were inspected for formation of a distinct clear zone due to cell killing.

### Antibacterial activity of campycins *in vitro*

To assess the killing efficiency of campycins under different conditions, *C. jejuni* strains were grown on BA plates and harvested with CBHI broth and adjusted to the final concentration of 10^6^ cfu/ml. 100 μl of each culture (10^6^ cfu/ml) was mixed with 100μl of each campycin (10^9^ campycins/ml) for 3 or 24 h under MA conditions or 24 h under anaerobic conditions using an anaerobic gas generator (AnaeroGen), in plastic containers at 5 °C. Cells were treated with TN50 instead of campycins as a negative control. After incubation, samples were tenfold diluted in TN50 buffer and spotted on BA plates that were incubated for 48 h under MA conditions. Colonies were enumerated and cfu/ml calculated. Data represent the mean from biological triplicates.

### Construction of plasmids for H-fiber expression

To express CJIE1 H-fibers of either CAMSA2147 or RM1221, the responsible genes were cloned into vector pET28-a (+) by using In-Fusion^®^ cloning – Takara Bio. Specifically, CAMSA2147 (GCA_003095855) was used as a template for amplification of the fragment comprising the coding sequence of the C-terminus of the H-fiber (amino acids 151 to 343) by specific primers (Table S3). The pET28-a (+) vector was digested with NdeI and XhoI and recombination resulted in the pCRYS1 plasmid. Similarly, RM1221 (CP000025) was used as a template for amplification of the fragment comprising the coding sequence of the C-terminus of the H-fiber (amino acids 151 to 359) with or without the downstream predicted chaperone gene. Specific primers were used for the amplification (Table S3) and recombination of the fragments with the linearized pET28-a (+) resulted in the pCRYS4 and pCRYS5 plasmids, respectively (Table S2). *E. coli* BL21-CodonPlus-(DE3)-RIL were used for transformation and transformants were selected on LB agar plates in the presence of kanamycin (100 μg/ml) and chloramphenicol (50 μg/ml).

H-fibers were expressed with or without the predicted chaperones by growing cells in 1-liter of LB at 37 °C until reaching optical density OD_600_ = 0.6. Protein expression was induced by adding isopropyl-beta-D-thiogalactopyranoside (0.5 mM) and cultures were incubated for 18 h at 16 °C at 120 rpm. Pellets of cultures were harvested by centrifugation (8,000 × *g*, 10 min, 4 °C) and resuspended in 10 ml of lysis buffer (20 mM NaH_2_PO_4_-NaOH, 0.5 M NaCl, 50 mM imidazole, pH 7.4). Sonication (Bandelin Sonopul HD 2070 homogeniser) with 10 bursts of 30 sec (amplitude of 50%) and 30 sec intervals allowed cell lysis and samples were further filtered twice with 0.22 μm pore size filters. His GraviTrap^TM^ gravity flow columns (GE Healthcare) were used for protein purification. The lysis buffer was used for the wash step during purification and proteins were eluted with 6 ml of elution buffer (20 mM NaH2PO4-NaOH, 0.5 M NaCl, 500 mM imidazole, pH 7.4). The elution buffer was further exchanged with Thermo Scientific BupH Tris Buffered Saline (TBS) by using Amicon Ultra-15 Centrifugal Filter Units with Ultracel-10 membrane cutoff (Merck Millipore) and protein concentration was measured with a Qubit 2.0 fluorometer.

### Determination of the H-fiber receptor

For identifying the receptor, we used the Pierce™ His Protein Interaction Pull-Down Kit that purifies protein interactors of any His-tagged fusion proteins. To do so, we used 700 μl of 200 μg purified H-fibers fused with a His-tag as a bait. First, the H-fibers were immobilized onto cobalt chelate resin and further trapped their prey that was present in the cell lysates. CAMSA2147 or RM1221 were used for providing the prey protein i.e., the receptor. Specifically, the strains were grown on BA plates and cells were harvested using a loop and dissolved in 1 ml of TBS. Cell lysates were further prepared by following the manufacturer’s instructions. Either the H-fibers or the cell lysates alone were run in the columns as negative controls. Finally, 30 μl of each eluted sample was analyzed by SDS-PAGE analysis and the bands of interest were cut from the gel for further analysis. Alphalyse A/S and Proteome Factory further identified the proteins using LCMS-based high resolution mass spectrometric techniques.

### Bioinformatic analysis

To investigate the diversity of CJIE1 H-fibers, CAMSA2147 H-fiber (WP_002878910.1) was used for searching CJIE1 H-fiber homologs in our CAMSA bacterial collection and sequences were further aligned using CLC Main Workbench 22 (QIAGEN). To analyse the MOMP diversity, CAMSA2147 MOMP (WP_002875862.1) was used as a query sequence to search *C. jejuni* homologous proteins in the National Center for Biotechnology Information (NCBI) genome database through BLASTP (34). In total 5.000 MOMP sequences were retrieved that were further aligned, including default settings in mafft (35) and neighbor-joining phylogenetic tree was created, using 100 bootstraps. Interactive Tree Of Life (iTOLv6) was used for the display, manipulation and annotation of the phylogenetic tree (36). Structural prediction of proteins and protein complexes was performed using Colab Alphafold2 (37) and visualized using PyMOL (38).

### Statistical analysis

GraphPad Prism 7 software (Version 7.0d) was used for statistical analysis. To test the significance of logarithmic bacterial reduction of cells treated with campycin compared to the cells treated with TN50 buffer, we used the Paired-Samples t-test with 95% confidence interval percentage based on biological triplicates.

## SUPPLEMENTAL MATERIAL

Supplementary figures and tables are provided as a single file and available online.

## DATA AVAILABILITY

Material is available upon request by contacting the corresponding author Martine Camilla Holst Sørensen. Accession numbers when appropriate are listed in the supplemental material.

## ACKNOWLEDGEMENT

The work was supported by the Danish AgriFish Agency of Ministry of Environment and Food (34009-14-0873) and the European Union’s horizon 2020 Marie Skladowska Curie-Individual Fellowship (705817). The salary of M.C.H.S was partially funded by Intralytix, Inc., which had no influence on the design or conclusions of the present work. We sincerely thank our colleague Anders Nørgaard Sørensen for valuable advice in the preparation of the manuscript and finalizing figures in PyMOL. The authors declare no competing interests. Conceptualization: A.Z., Y.E.G., L.B., M.C.H.S.; methodology: A.Z., Y.E.G., L.B., M.C.H.S.; investigation: A.Z., Y.E.G.; formal analysis: A.Z., Y.E.G.; writing – original draft: A.Z., M.C.H.S.; writing – final draft: A.Z., Y.E.G., L.B., M.C.H.S.; funding acquisition: Y.E.G., L.B., M.C.H.S.; resources, M.C.H.S., L.B.; supervision, L.B., M.C.H.S.

## REFERENCES

1. Williams SR, Gebhart D, Martin DW, Scholl D. 2008. Retargeting R-type pyocins to generate novel bactericidal protein complexes. Appl Environ Microbiol 74:3868–3876. 10.1128/aem.00141-08.

2. Michel-Briand Y, Baysse C. 2002. The pyocins of *Pseudomonas aeruginosa*. Biochimie 84:499–510. 10.1016/s0300-9084(02)01422-0.

3. Dams D, Brøndsted L, Drulis-Kawa Z, Briers, Y. 2019. Engineering of receptor-binding proteins in bacteriophages and phage tail-like bacteriocins. Biochem Soc Trans, 47:449–460. 10.1042/bst20180172.

4. Zampara A, Sørensen MCH, Grimon D, Antenucci F, Vitt AR, Bortolaia V, Briers Y, Brøndsted L. 2020. Exploiting phage receptor binding proteins to enable endolysins to kill Gram-negative bacteria. Sci Rep 10:12087. 10.1038/s41598-020-68983-3.

5. Scholl D, Cooley M, Williams SR, Gebhart D, Martin D, Bates A, Mandrell R. 2009. An engineered R-type pyocin is a highly specific and sensitive bactericidal agent for the food-borne pathogen *Escherichia coli* O157:H7. Antimicrob Agents Chemother 53:3074–3080. 10.1128/aac.01660-08.

6. Hermans D, Van Deun K, Martel A, Van Immerseel F, Messens W, Heyndrickx M, Haesebrouck F, Pasmans F. 2011. Colonization factors of *Campylobacter jejuni* in the chicken gut. Vet Res 42:82. 10.1186/1297-9716-42-82.

7. Igwaran A, Okoh AI. (2019). Human campylobacteriosis: A public health concern of global importance. Heliyon 5:e02814. 10.1016/j.heliyon.2019.e02814.

8. Devleesschauwer B, Bouwknegt M, Mangen MJJ, Havelaar AH. 2017. Health and economic burden of Campylobacter, p 27-40. In Klein G (ed), Campylobacter, Academic Press, Elsevier.

9. Clark CG, Ng LK. 2008. Sequence variability of *Campylobacter* temperate bacteriophages. BMC Microbiol 8:49. 10.1186/1471-2180-8-49.

10. Zampara A, Sørensen MCH, Gencay YE, Grimon D, Kristiansen SH, Jørgensen LS, Kristensen JR, Briers Y, Elsser-Gravesen A, Brøndsted L. 2021. Developing innolysins against *Campylobacter jejuni* using a novel prophage receptor-binding protein. Front Microbiol 12:619028. 10.3389/fmicb.2021.619028.

11. Sørensen MCH, Gencay YE, Birk T, Baldvinsson SB, Jäckel C, Hammerl JA, Vegge CS, Neve H, Brøndsted L. 2015. Primary isolation strain determines both phage type and receptors recognised by *Campylobacter jejuni* bacteriophages. PloS One 10:e0116287. 10.1371/journal.pone.0116287.

12. Tanoeiro L, Oleastro M, Nunes A, Marques AT, Duarte SV, Gomes JP, Matos APA, Vítor JMB, Vale FF. 2022. Cryptic prophages contribution for *Campylobacter jejuni* and *Campylobacter coli* introgression. Microorganisms 10:516. 10.3390/microorganisms10030516.

13. Ostenfeld LJ, Sørensen AN, Neve H, Vitt A, Klumpp J, Sørensen MCH. 2023. A hybrid receptor binding protein enables phage F341 infection of *Campylobacter* by binding to flagella and lipooligosaccharides. bioRxiv 10.1101/2023.09.05.556331.

14. Haggard-Ljungquist E, Halling C, Calendar R. 1992. DNA sequences of the tail fiber genes of bacteriophage P2: evidence for horizontal transfer of tail fiber genes among unrelated bacteriophages. J Bacteriol 174:1462–1477. 10.1128/jb.174.5.1462-1477.1992.

15. North OI, Sakai K, Yamashita E, Nakagawa A, Iwazaki T, Büttner CR, Takeda S, Davidson AR. 2019. Phage tail fibre assembly proteins employ a modular structure to drive the correct folding of diverse fibres. Nat Microbiol 4:1645–1653. 10.1038/s41564-019-0477-7.

16. Djordjevic SP, Unicomb LE, Adamson PJ, Mickan L, Rios R. 2007. Clonal complexes of *Campylobacter jejuni* identified by multilocus sequence typing are reliably predicted by restriction fragment length polymorphism analyses of the *flaA* gene. J Clin Microbiol 45:102–108. 10.1128/jcm.01012-06.

17. Wirz SE, Overesch G, Kuhnert P, Korczak BM. 2010. Genotype and antibiotic resistance analyses of *Campylobacter* isolates from ceca and carcasses of slaughtered broiler flocks. Appl Environ Microbiol 76:6377–6386. 10.1128/aem.00813-10.

18. Ferrara LGM, Wallat GD, Moynié L, Dhanasekar NN, Aliouane S, Acosta-Gutiérrez S, Pagès JM, Bolla JM, Winterhalter M, Ceccarelli M, Naismith JH. 2016. MOMP from *Campylobacter jejuni* is a trimer of 18-Stranded β-barrel monomers with a Ca^2+^ Ion Bound at the Constriction Zone. J Mol Biol 428:4528–4543. doi: 10.1016/j.jmb.2016.09.021. https://doi.org/10.1016/j.jmb.2016.09.021.

19. Zhang Q, Meitzler JC, Huang S, Morishita T. 2000. Sequence polymorphism, predicted secondary structures, and surface-exposed conformational epitopes of *Campylobacter major* outer membrane protein. Infect Immun 68:5679–5689. 10.1128/iai.68.10.5679-5689.2000.

20. Taha-Abdelaziz K, Singh M, Sharif S, Sharma S, Kulkarni RR, Alizadeh M, Yitbarek A, Helmy YA. 2023. Intervention strategies to control *Campylobacter* at different stages of the food chain. Microorganisms 11:113.

21. Klumpp J, Dunne M, Loessner MJ. 2023. A perfect fit: Bacteriophage receptor-binding proteins for diagnostic and therapeutic applications. Curr Opin Microbiol 71:102240. 10.1016/j.mib.2022.102240.

22. Gonzalez-Serrano R, Rosselli R, Roda-Garcia JJ, Martin-Cuadrado AB, Rodriguez-Valera F, Dunne M. 2023. Distantly related *Alteromonas* bacteriophages share tail fibers exhibiting properties of transient chaperone caps. Nat Commun 14:6517. 10.1038/s41467-023-42114-8.

23. Cody AJ, Maiden MJC, Dingle KE. 2009. Genetic diversity and stability of the *porA* allele as a genetic marker in human *Campylobacter* infection. Microbiology (Reading) 155:4145–4154. 10.1099/mic.0.031047-0.

24. Colles FM, Preston SG, Barfod KK, Flammer PG, Maiden MCJ, Smith AL. 2019 Parallel sequencing of porA reveals a complex pattern of *Campylobacter* genotypes that differs between broiler and broiler breeder chickens. Sci Rep. 9:6204. 10.1038/s41598-019-42207-9.

25. Kropinski AM, Lewis V, Berry D. 1987. Effect of growth temperature on the lipids, outer membrane proteins, and lipopolysaccharides of *Pseudomonas aeruginosa* PAO. J Bacteriol 169:1960–1966. doi:10.1128/jb.169.5.1960-1966.1987.

26. Burnham PM, Hendrixson DR. 2018. *Campylobacter jejuni*: collective components promoting a successful enteric lifestyle. Nat Rev Microbiol. 16:551–565. 10.1038/s41579-018-0037-9.

27. EFSA BIOHAZ Panel (EFSA Panel on Biological Hazards), Koutsoumanis, K, Allende A, Alvarez-Ordóñez A, Bolton D, Bover-Cid S, Davies R, De Cesare A, Herman L, Hilbert F, Lindqvist R, Nauta M, Peixe L, Ru G, Simmons M, Skandamis P, Suffredini E, Alter T, Crotta M, Ellis-Iversen J, Hempen M, Messens W, Chemaly M. 2020. Update and review of control options for *Campylobacter* in broilers at primary production. EFSA Journal 2020 18:6090. 10.2903/j.efsa.2020.6090.

28. Gebhart D, Lok S, Clare S, Tomas M, Stares M, Scholl D, Donskey CJ, Lawley TD, Govoni GR. 2015. A modified R-type bacteriocin specifically targeting *Clostridium difficile* prevents colonization of mice without affecting gut microbiota diversity. mBio 6:e02368–14. 10.1128/mbio.02368-14.

29. Ritchie JM, Greenwich JL, Davis BM, Bronson RT, Gebhart D, Williams SR, Martin D, Scholl D, Waldor MK. 2011. An *Escherichia coli* O157-specific engineered pyocin prevents and ameliorates infection by *E. coli* O157:H7 in an animal model of diarrheal disease. Antimicrob Agents Chemother 55:5469–74. 10.1128/aac.05031-11.

30. Choi KH, Kumar A, Schweizer HP. 2006. A 10-min method for preparation of highly electrocompetent *Pseudomonas aeruginosa* cells: Application for DNA fragment transfer between chromosomes and plasmid transformation. J Microbiol Methods 64:391–397. 10.1016/j.mimet.2005.06.001.

31. Ikeda K, Egami F. 1969. Receptor substance for pyocin R. I. Partial purification and chemical properties. J Biochem 65:603–609. 10.1093/oxfordjournals.jbchem.a129053.

32. Kageyama M, Egami F. 1962. On the purification and some properties of a pyocin, a bacteriocin produced by *Pseudomonas aeruginosa*. Life Sci 1:471–6. 10.1016/0024-3205(62)90055-3.

33. Gencay YE, Birk T, Sørensen MCH, Brøndsted L. 2017. Methods for isolation, purification, and propagation of bacteriophages of *Campylobacter jejuni*. Methods Mol Biol 1512:19–28. 10.1007/978-1-4939-6536-6_3.

34. Sayers EW, Bolton EE, Brister JR, Canese K, Chan J, Comeau DC, Connor R, Funk K, Kelly C, Kim S, Madej T, Marchler-Bauer A, Lanczycki C, Lathrop S, Lu Z, Thibaud-Nissen F, Murphy T, Phan L, Skripchenko Y, Tse T, Wang J, Williams R, Trawick BW, Pruitt KD, Sherry ST. 2022. Database resources of the national center for biotechnology information. Nucleic Acids Res 50:D20–D26. 10.1093/nar/gkab1112.

35. Katoh K, Rozewicki J, Yamada KD. 2019. MAFFT online service: multiple sequence alignment, interactive sequence choice and visualization. Briefings in bioinformatics 20:1160–1166.

36. Letunic I, Bork P. 2021. Interactive Tree Of Life (iTOL) v5: an online tool for phylogenetic tree display and annotation. Nucleic Acids Res 49:W293–W296. 10.1093/nar/gkab301.

37. Mirdita M, Schütze K, Moriwaki Y, Heo L, Ovchinnikov S, Steinegger M. 2022. ColabFold: making protein folding accessible to all. Nat Methods 19:679–682. 10.1038/s41592-022-01488-1.

38. Schrodinger LLC. 2015. The PyMOL Molecular Graphics System, Version 1.8.

